# Auxin-independent effects of apical dominance induce temporal changes in phytohormones

**DOI:** 10.1101/2022.10.24.513542

**Authors:** Da Cao, Francois Barbier, Elizabeth A. Dun, Franziska Fichtner, Lili Dong, Stephanie C. Kerr, Christine A. Beveridge

## Abstract

The inhibition of shoot branching by the growing shoot tip of plants, termed apical dominance, was originally thought to be mediated by auxin. Recently the importance of the shoot tip sink strength during apical dominance has re-emerged with recent studies highlighting roles for sugars in promoting branching. This raises many unanswered questions on the relative roles of auxin and sugars in apical dominance. Here we show that auxin regulation of cytokinins, which promote branching, is significant only after an initial stage of branching we call bud release. During this early bud release stage, rapid cytokinin increases are associated with enhanced sugars. Auxin may also act through strigolactones which have been shown to suppress branching after decapitation, but here we show that strigolactones do not have a significant effect on initial bud outgrowth after decapitation. We report here that when sucrose or cytokinin is abundant, strigolactones are less inhibitory during the bud release stage compared to later stages and that strigolactone treatment rapidly inhibits cytokinin accumulation in pea axillary buds of intact plants. After initial bud release, we find an important role of gibberellin in promoting sustained bud growth downstream of auxin. We are therefore able to suggest a model of apical dominance that integrates auxin, sucrose, strigolactones, cytokinins and gibberellins and describes differences in signalling across stages of bud release to sustained growth.

## Introduction

Shoot branching is an important determinant of shoot architecture and affects the yield and/or value of most agricultural, forestry and ornamental crops. Apical dominance is a form of branching control whereby the growing shoot tip inhibits the outgrowth of axillary buds (Phillips, 1975; Ongaro and Leyser, 2007; Barbier et al., 2017). Under apical dominance, removal of the shoot tip by herbivory, pruning or decapitation releases axillary buds to grow.

Since the pioneering work of Sachs and Thimann (1964), plant hormones have been proposed as key mediators of apical dominance whereby auxin produced in the main shoot tip is transported downwards and indirectly inhibits axillary bud growth. Auxin reduces the supply of the stimulatory hormone cytokinin (CK) to axillary buds through suppressing CK content in stems and this concept is widespread and observed across diverse plants (Nordström et al., 2004; Tanaka et al., 2006; Su et al., 2011). Auxin also enhances the expression of strigolactone (SL) biosynthesis genes which is thought to enhance the supply of this bud growth inhibitor to buds (Sorefan et al., 2003; Foo et al., 2005; Hayward et al., 2009). Application of CK directly on axillary buds triggers their outgrowth, while application of SL represses their outgrowth (Gomez-Roldan et al., 2008; Brewer et al., 2009; Dun et al., 2012; Tan et al., 2019). In many species, CK and SL largely act antagonistically through a common transcription factor TEOSINTE BRANCHED 1 (BRC1) which is the gene largely responsible for the difference in branching between the non-tillering maize and its high tillering wild progenitor teosinte. Expression of *BRC1* is correlated with bud inhibition (Braun et al., 2012; Dun et al., 2012, 2013; Seale et al., 2017; Kerr et al., 2020), and *brc1* deficient mutants show an increased branching phenotype in several plant species from divergent groups of angiosperms (Aguilar-Martínez et al., 2007; Martín - Trillo et al., 2011; Ramsay et al., 2011; Studer et al., 2011; Braun et al., 2012).

Interactions between SL and CK pathways have been reported (Dun et al., 2012; Duan et al., 2019; Kerr et al., 2021). CK rapidly regulates transcript levels of DWARF53 (D53), which is a negative regulator of SL signalling in rice, and its homologues *SMXL6/7/8* in pea (Kerr et al., 2021). In rice, SL can promote CK degradation through transcriptionally enhancing *CYTOKININ OXIDASE 9 (OsCKX9*) (Duan et al., 2019). This finding is supported by higher CK content in SL deficient mutant shoots compared with wild type (WT) plants in pea and rice (Young et al., 2014; Duan et al., 2019). However, exogenous SL supply in rice reduces bud outgrowth but does not affect CK levels or the expression of CK biosynthesis genes in tiller nodes (Xu et al., 2015).

Auxin movement in the polar auxin transport stream may suppress branching through the competition of auxin flow between main stem and axillary bud (Prusinkiewicz et al., 2009; Balla et al., 2011; Balla et al., 2016). This model has been prominent in arabidopsis where bud inhibition has been associated with inhibition of auxin transport from buds, relative to auxin flow in the main stem. This correlation has not held up in terms of the initial growth of pea buds after decapitation or CK treatment (Brewer et al., 2015; Chabikwa et al., 2019). Instead, reduced auxin transport specifically in buds (and not stem) had no growth inhibition effect for two days after the induction of bud growth (Brewer et al., 2015; Chabikwa et al., 2019). This early stage of bud growth has not generally been explored in other model systems and hence it is not clear as to whether auxin transport is involved in the induction of bud release in diverse plants or, as in pea, it may be more relevant at advanced stages of bud outgrowth.

Changes in auxin level and transport in buds relative to stem have been proposed to promote branching (Gocal et al., 1991; Leyser, 2006; Barbier et al., 2015; Leyser, 2018). Several studies using garden pea have questioned this model finding no correlation between auxin transport from buds and their early growth (bud release)(Brewer et al., 2015; Chabikwa et al., 2019). Here we question whether another means through which auxin in buds may affect bud outgrowth is through the well-established role of auxin in regulating gibberellin (GA) levels (Scott et al., 1967; Ross et al., 2000; Wolbang and Ross, 2001; Ross et al., 2003; Zhu et al., 2022). A stimulatory role of GA in bud growth has been widely reported in tree species (Elfving et al., 2011; Ni et al., 2015; Tan et al., 2018; Katyayini et al., 2020) but less so for herbaceous plants and grass species (Kebrom et al., 2013). Exogenous treatment of GA to buds can break bud dormancy in the woody plant *Jatropha curcas*, potentially via inhibiting the expression of *BRC1* (Ni et al., 2015). Locally increased GA biosynthesis gene expression in buds of the *brc1* mutant in maize also indicates that BRC1 may inhibit GA production and signalling (Dong et al., 2019). This warrants testing of the hypothesis that rising auxin levels in buds may regulate bud GA levels to specifically promote bud growth.

Several studies have associated dwarfism with increased branching including across a range of lines affected in GA level or response (Sasaki et al., 2002; Lo et al., 2008; Liao et al., 2019). As discussed below, additional resources available for axillary buds due to suppressed main stem growth in dwarf plants, could enhance shoot branching. Similarly, given the rapid growth response of the shoot tip of many herbaceous plants in response to exogenous GA, it is difficult to interpret the branching response of many GA treatment experiments due to competition between buds and main shoot growth.

The demand of the shoot tip for sugars has recently re-emerged as an important component of apical dominance (Barbier et al., 2015; Kebrom, 2017; Schneider et al., 2019; Kotov et al., 2021). This renewed attention on sugars including sucrose, the mobile product of photosynthesis, is partly because the dynamics of auxin depletion after decapitation are too slow to account for initial bud outgrowth whereas changes in sucrose are rapid (Morris et al., 2005; Mason et al., 2014). Axillary bud outgrowth is promoted by sugars in different plant species (Mason et al., 2014; Barbier et al., 2015; Xia et al., 2021) and the enhanced supply of sugars after decapitation is sufficiently rapid to correlate with the timing of bud release (Mason et al., 2014). Levels of trehalose 6-phosphate (Tre6P), a low abundant metabolite that signals sucrose availability (Fichtner and Lunn, 2021), also increase in axillary buds after decapitation and this increase is correlated with the onset of bud outgrowth (Fichtner et al., 2017). The branching phenotypes of arabidopsis mutants with altered levels of Tre6P (Yadav et al., 2014; Fichtner et al., 2021) or altered glucose signalling via HEXOKINASE 1 (Barbier et al., 2021) support a signalling role of sugars in regulation of shoot branching (Barbier et al., 2019).

One effect of sugars in the regulation of shoot branching may be to promote CK accumulation and suppress SL signalling. In experiments examining bud growth *in vitro*, sucrose treatment increased CK levels in nodal stems of rose and dark-grown stems of potato and promoted bud outgrowth (Barbier et al., 2015; Salam et al., 2021). In etiolated (dark-grown) potato sprouts, sugars are very important for bud outgrowth. Sucrose feeding increases CK production and exogenous CK can promote bud growth in etiolated potato sprouts even without exogenous sucrose supply. The possibility that sucrose rapidly promotes of cytokinin levels in decapitated plants has not been tested in vivo in separation from auxin depletion.

Sucrose can repress SL inhibition of bud growth in a variety of plant species (Dierck et al., 2016; Bertheloot et al., 2020; Patil et al., 2021). Studies show that sucrose may be involved in reducing SL signalling as sucrose can repress expression of the SL signalling gene, *DWARF3*, and promote accumulation of D53 in rice and pea.

The possibility that sugars may directly and rapidly affect CK levels and SL signalling is far removed from the initial model of auxin-mediated apical dominance. This paradigm shift in apical dominance thinking is yet to be tested on light-grown plants with manipulations of apical dominance *in vivo*. In this study we address this by investigating responses of buds in decapitated plants and in relation to timing of changes in auxin content.

## Results

Previous studies in pea showed that auxin depletion in internodes close to the site of decapitation can regulate local CK levels (Tanaka et al., 2006). In this study, we used tall plants with additional internodes (Figure 1) such that the zone of auxin depletion in the main stem remained above node 4 close to the site of decapitation (upper region), but did not extend to node 2 (lower region) at 6 h after decapitation (Figure 1C). The upper region was useful to repeat the widely observed correlation of auxin depletion with enhanced CK levels whilst the lower region (at and just above node 2) served to explore the phytohormone properties associated with bud outgrowth outside the zone of main stem auxin depletion. After decapitation in these plants, significant bud growth (2 h; Figure 1A) and reduced *BRC1* gene expression (3h; Figure 1B) was observed outside the zone of stem auxin depletion as reported previously (Mason et al., 2015).

**Figure 1.**
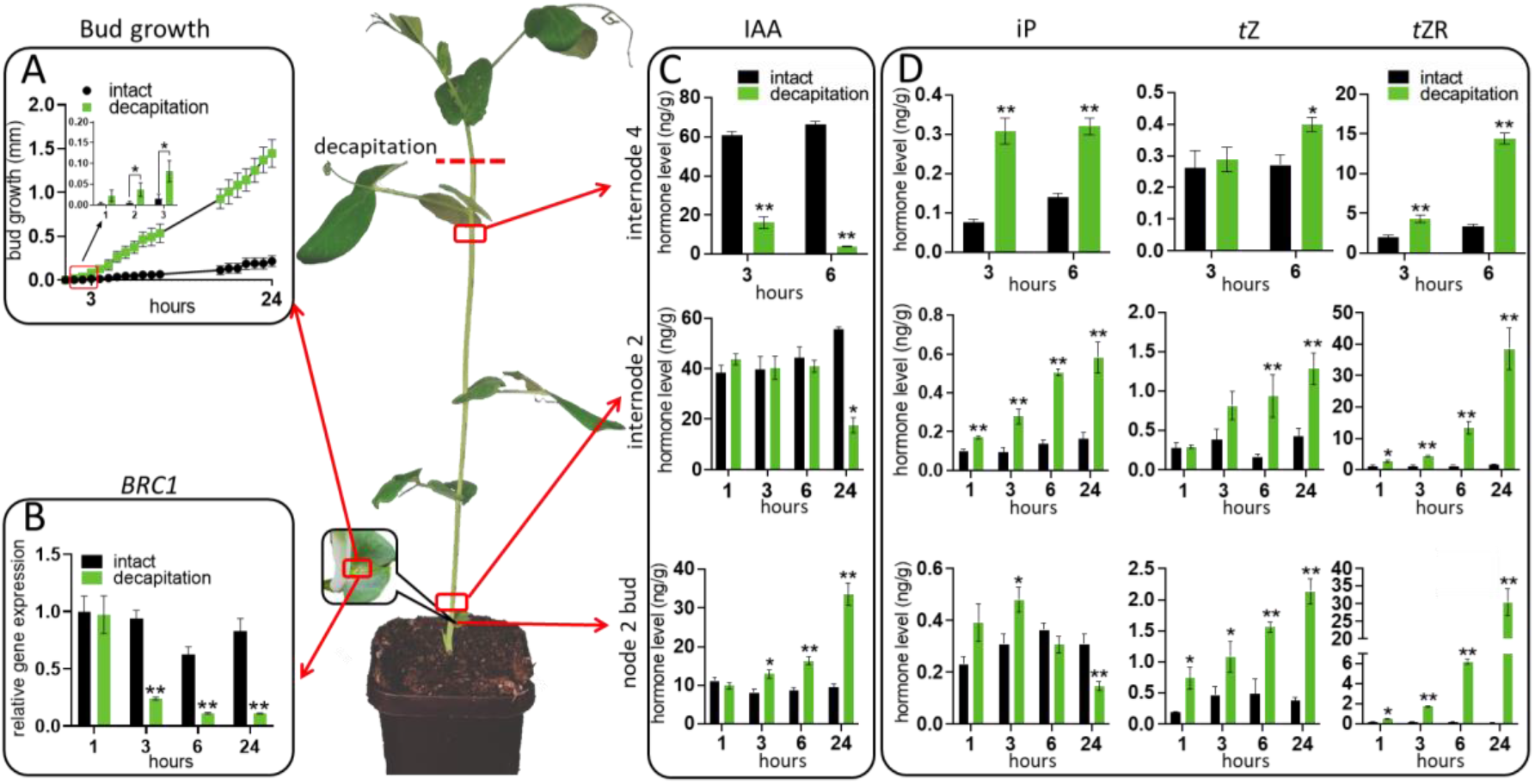
Tall plants enabled an exploration of growth (A), gene expression (B) and changes in hormone level (C, D) in buds within and below a zone of auxin depletion. (A) Growth of node 2 wild-type buds in decapitated and intact plants. *P* <0.05, one-tailed Student’s *t* test, *n* = 5. (B) Expression of *BRC1* in node 2 buds at 1, 3, 6, and 24 h after decapitation. Results are presented relative to intact control at 1 h, *n* = 4. (C, D) Endogenous auxin (C) and cytokinin levels (D) in internode 2, internode 4 and node 2 bud at 1, 3, 6, and 24 h after decapitation, *n* = 4. Each replicate contains 20 individual buds. Node 2 was approximately 12 cm from the decapitation site. Values are mean ± SE. * *P*<0.05, \*\**P* < 0.01, two-tailed Student’s *t* test for B, C and D. IAA, indole-3-acetic acid; iP, isopentenyladenine; *t*Z, *trans*-zeatin; *t*ZR, *trans*-zeatin riboside.

### Endogenous CK levels increase before measurable bud growth and are not correlated with auxin depletion

To investigate whether changes in CK levels occurred outside the zone of auxin depletion, we quantified CK levels in internode 2 and 4 stem segments and node 2 buds in intact and decapitated plants (Figure 1D, and Supplemental Figure S1). The quantified CKs include three types of bioactive forms, isopentenyladenine (iP), *trans*-zeatin (*tZ*) and dihydrozeatin (DZ), and their precursors and transported forms including CK ribosides, which may also be bioactive (Nguyen et al., 2021), and CK nucleosides. As previously reported with smaller seedlings of pea (Tanaka et al., 2006), auxin levels reduced and levels of iP- and *t*Z-type of CKs increased in the upper stem, verifying the expected anti-correlation of auxin and CK within the zone of auxin depletion (internode 4 segment; Figure 1D). Auxin levels were not depleted in the stem at internode 2 until after 6 h. However, in this region outside the zone of auxin depletion, CK levels also increased rapidly in the stem (Figure 1D). In internode 2, all CKs except for *trans*-zeatin riboside-5’-monophosphate (*t*ZMP), *tZ* and DZ increased significantly at 1 h after decapitation. CK levels also increased in node 2 buds within 1 h and this included all types of CKs except for iP and DZ-riboside (Figure 1D, and Supplemental Figure S1). In node 2 buds, this significant increase in CKs was sustained or enhanced throughout the 24 h time course except for iP which first significantly increased at 3 h in buds and then stopped accumulating and showed a significant decrease at 24 h after decapitation relative to the intact control (Figure 1D). Interestingly, outside the zone of auxin depletion in the stem, the auxin content in axillary buds at node 2 increased significantly at 3 h and continued to rise thereafter as previously described in bean (Gocal et al., 1991).

To test whether auxin depletion close to the site of decapitation somehow indirectly triggers the distal increase in CK outside the zone of auxin depletion, we monitored changes in CK level and related gene expression in internode 2 and internode 4 after decapitation and treatment with or without the synthetic auxin 1-naphthaleneacetic acid (NAA) applied to the decapitated stump. As expected, NAA treatment was clearly absorbed (Supplemental Figure S2A) and effectively prevented decapitation-induced accumulation of CK nucleotides and CK ribosides and the expression of CK biosynthesis genes *ISOPENTYL TRANSFERASE1 (IPT1*) and *IPT2* (Figure 2A and 2C). This is consistent with previous findings in excised pea segments (Tanaka et al., 2006). In contrast, the accumulation of iP, *t*Z and DZ in internode 4 following decapitation was not reduced by exogenous auxin supply (Figure 2A). In fact, exogenous auxin supply to the decapitated stump unexpectedly increased the accumulation of these bioactive CKs at this 4 h time point. This is in line with auxin-boosted gene expression of *LONELY GUY1 (LOG1), LOG3* and *LOG7*, which catalyze the synthesis of bioactive CKs from CK nucleotides (Figure 2B). Coupled with auxin-induced decreased nucleotide levels, this is consistent with reduced overall CK levels as expected in the longer term (Tanaka et al., 2006; Young et al., 2014).

**Figure 2.**
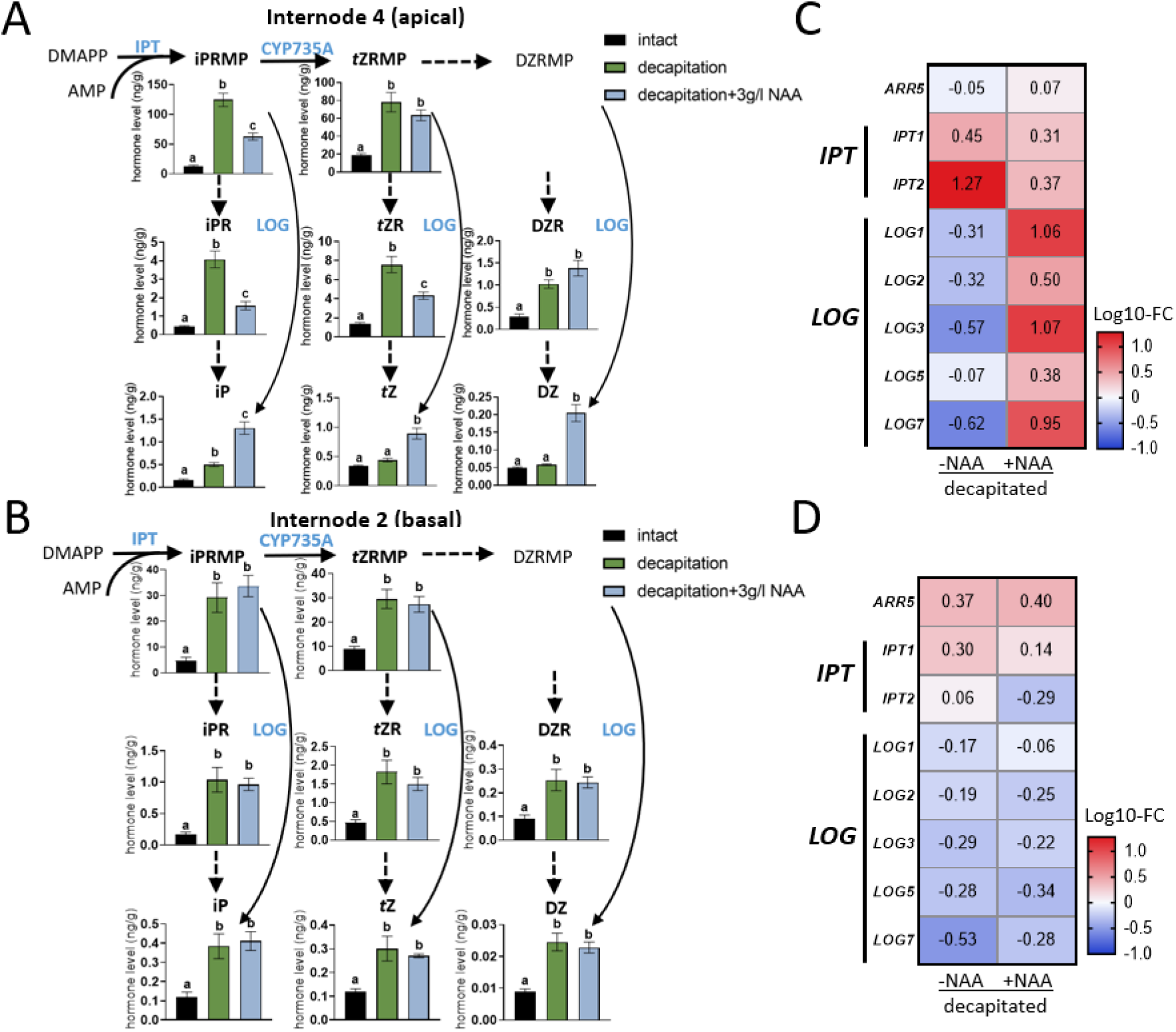
Decapitation-induced CK accumulation is not initially caused by auxin depletion. (A, B) Endogenous CK levels in internode 4 (A) and internode 2 (B) 4 h after decapitation. Decapitated shoots were treated either with mock or 3 g/L NAA above internode 4. Values are mean ± SE, *n* = 4. Multiple comparison tests were performed with one-way ANOVA. Different letters on the top of columns indicate statistically significant differences. (C, D) Log_10_ fold changes compared with intact plants in transcript of CK biosynthesis and signalling genes in internode 4 (C) and internode 2 (D) of decapitated plants treated either with mock or 3 g/L NAA above internode 4. Numbers represent fold change compared with intact plants. Abbreviations: DMAPP, dimethylallyl diphosphate; iPRMP, isopentenyladenosine-5’-monophosphate; *t*ZRMP, *trans*-zeatin riboside-5’-monophosphate; DZRMP, dihydrozeatin riboside-5’-monophosphate; iPR, isopentenyladenosine; *t*ZR, *trans*-zeatin riboside; DZR, dihydrozeatin riboside; iP, isopentenyladenine; *t*Z, *trans*-zeatin; DZ, dihydrozeatin; IPT, adenosine phosphate-isopentenyltransferase; LOG, cytokinin phosphoribohydrolase ‘Lonely guy’; CYP735A, cytochrome P450 mono-oxygenase; *ARR5*, type-A response regulator 5.

NAA did not move to internode 2 within 4 h (Supplemental Figure 2B) and did not significantly prevent decapitation-induced accumulation of any CK types and did not affect CK biosynthesis gene expression (Figure 2B and 2D) in this region. Together, these results indicate that the rapid accumulation of CK in internode 2 is unlikely caused by decapitation-induced auxin depletion.

### Sugar availability enhances CK levels in buds

Given that enhanced CK content in the lower stem region was not associated with depleted auxin, we hypothesised that decapitation-induced sucrose accumulation may be involved (Mason et al., 2014; Fichtner et al., 2017; Salam et al., 2021; Wang et al., 2021). To determine if elevated sugar levels might be able to enhance CK levels in pea, we measured endogenous CK levels in buds exposed to varied sugar availability. Buds of excised stem segments showed significant growth at 4 h when exposed to 50 mM sucrose (Figure 3A, Supplemental Figure S3). Indeed, buds of excised stem segments grown on 50 mM sucrose contained substantially increased CK levels at 3 h compared with buds grown on mannitol (osmotic control, Figure 3B). Treatment of CK at a concentration that stimulates bud growth in intact pea (Dun et al., 2012), 50 μM 6-benzylaminopurine (BA), could not significantly promote the outgrowth of excised buds if sucrose was not supplied (Figure 3C). BA enhanced bud outgrowth when sucrose was in the range of 2 to 25 mM, but had little additional effect at 50 mM sucrose (Figure 3C).

**Figure 3.**
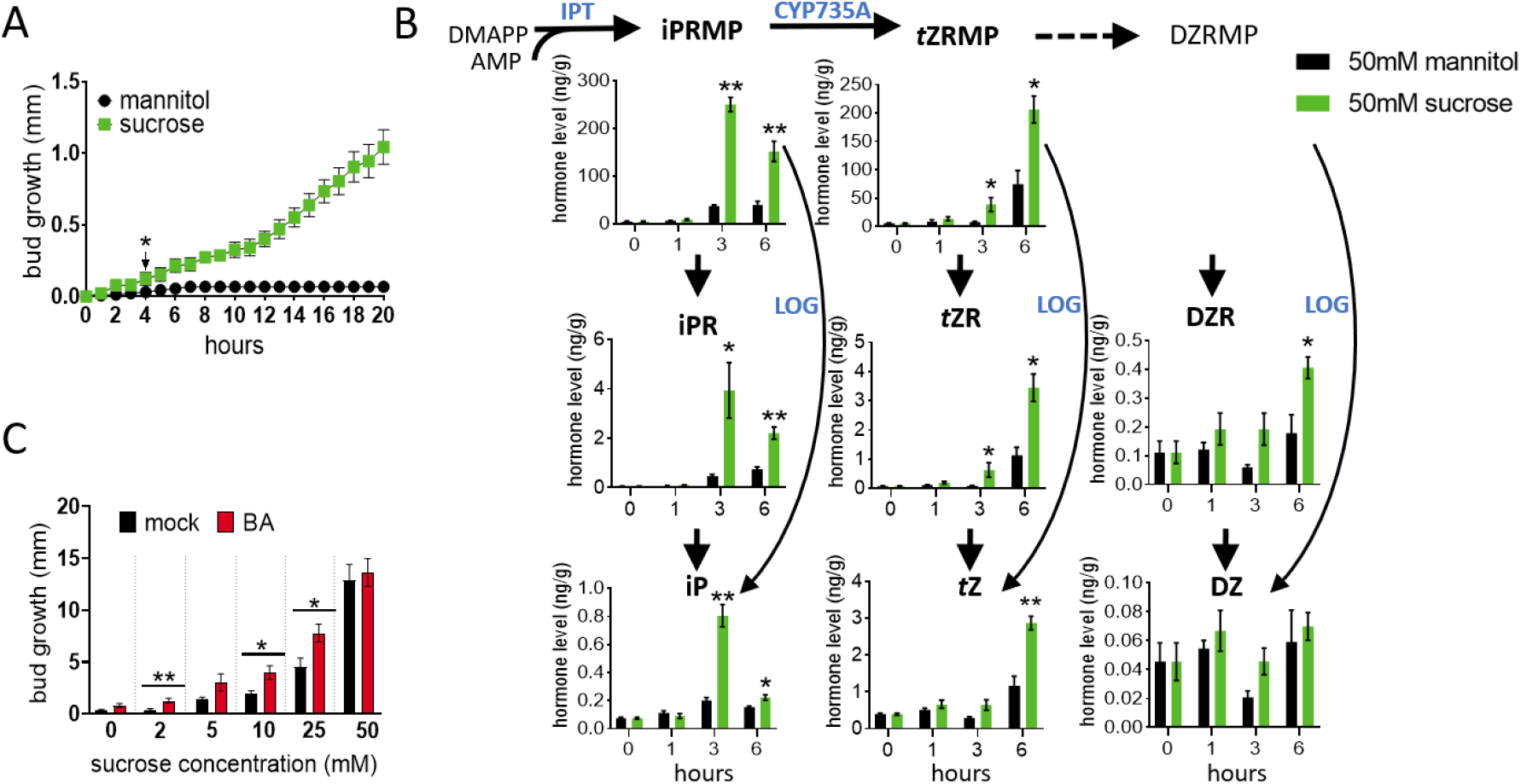
Sucrose initiates bud release and promotes CK accumulation in buds. (A) Outgrowth of buds on excised stem segments incubated with 50 mM sucrose or mannitol for 24 h, *n* = 5. * indicates timing of first significant difference, one-tailed Student’s *t* test. (B) Levels of endogenous CKs in buds on excised stem segments incubated with 50 mM sucrose or mannitol, *n* = 4. Each replicate contains 20 individual buds. Values are mean ± SE. * P<0.05, **P< 0.01, two-tailed Student’s *t* test. (C) Outgrowth of buds on excised stem segments incubated with 0, 2, 5,10, 25, 50 mM sucrose and treated with or without 50 μM 6-benzylaminopurine (BA) for 5 days, *n* = 5. Values are mean ± SE. * P<0.05, **P< 0.01, two-tailed Student’s *t* test. DMAPP, dimethylallyl diphosphate; iPRMP, isopentenyladenosine-5’-monophosphate; *t*ZRMP, *trans*-zeatin riboside-5’-monophosphate; DZRMP, dihydrozeatin riboside-5’-monophosphate; iPR, isopentenyladenosine; *t*ZR, *trans*-zeatin riboside; DZR, dihydrozeatin riboside; iP, isopentenyladenine; *t*Z, *trans*-zeatin; DZ, dihydrozeatin; IPT, adenosine phosphate-isopentenyltransferase; LOG, cytokinin phosphoribohydrolase ‘Lonely guy’; CYP735A, cytochrome P450 mono-oxygenases.

### SL reduces CK content in buds

To further study the interconnectivity among signals regulating shoot branching, we explored the effect of SL treatment on CK levels in axillary buds. GR24 (synthetic SL analogue) treatment to *ramosus5 (rms5*) SL-deficient mutant buds strongly inhibited bud outgrowth (Figure 4A) and reduced endogenous CK levels in the buds within 6 h after treatment (Figure 4B, Supplemental Figure S4A). To determine if this was due to a local effect of GR24 on CK levels in the bud, we also profiled CK levels in adjacent stem tissues at the same time point and found no change (Supplemental Figure S4A).

**Figure 4.**
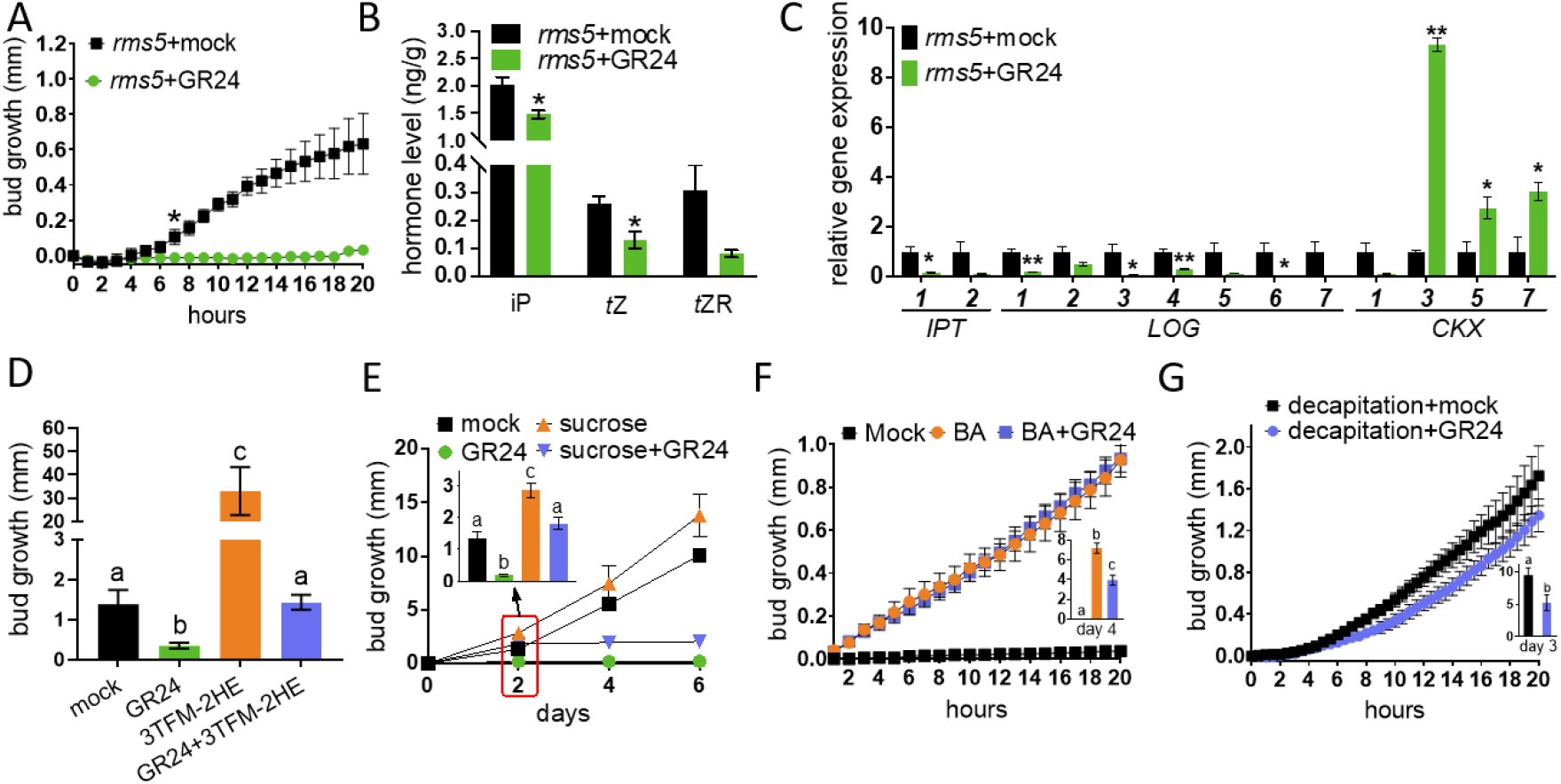
SL acts antagonistically with CK and sugars to inhibit axillary bud outgrowth and reduces CK levels in buds. (A) Growth of node 2 buds of *rms5* mutants treated with or without 1 μM GR24 (a synthetic SL). * indicates timing of first significant difference. One-tailed Student’s *t* test; *n* = 4. (B) Endogenous CK levels in *rms5* node 2 buds treated with or without 10 μM of GR24 for 6 h. *n* = 3. Each replicate contains 20 individual buds. (C) Expression levels of CK metabolism genes in *rms5* node 2 buds treated with or without 10 μM of GR24 for 6 h. *n* =3. Each replicate contains 20 individual buds. (D) Growth of *rms5* node 4 buds treated with mock, 10 μM GR24, 100 μM 3TFM-2HE, or 10 μM GR24 with 100 μM 3TFM-2HE (a CK oxidase inhibitor) after 7 days; *n* = 7. (E) Node 2 buds of *rms1* mutants treated with or without 5 μM GR24 and with or without 600 mM sucrose supplied to the stem vasculature. *n* = 4-6. (F) Growth of wild-type node 2 buds treated with or without 5 μM GR24 and 50 μM BA. *n* = 4-6. (G) Growth of wild-type node 2 buds treated with or without 1 μM GR24 and with decapitation at internode 8. *n* = 4-10. *rms5* plants with four fully expanded leaves were used for A-D. All values are mean ± SE. * P <0.05; ** P < 0.01 compared with mock control, two-tailed Student’s *t* test for B, C and G. One-way ANOVA for D, E, and F. Different letters on the top of columns indicate statistically significant differences. iP, isopentenyladenine; *t*Z, *trans*-zeatin; *t*ZR, *trans*-zeatin riboside.

To gain insight into the cause of decreased CK in buds after SL treatment, we quantified the expression of genes encoding CK biosynthesis and metabolism enzymes (Figure 4C) (Dun et al., 2012; Dolgikh et al., 2017). GR24 treatment on *rms5* buds not only significantly increased CK catabolism by promoting the gene expression of *CKXs (CKX3, CKX5* and *CKX7*), but also strongly inhibited CK biosynthesis by inhibiting the expression of the two *IPT* genes and five of the *LOG* genes (Figure 4C). In addition, GR24 treatment significantly increased the expression of bud dormancy marker genes*, DORMANCY-ASSOCIATED PROTEIN1 (DRM1*) and *BRC1* at 6 h after treatment (Supplemental Figure S4B). These results demonstrate that SL may inhibit CK levels in pea buds by decreasing CK biosynthesis and increasing CK degradation.

To investigate whether increased endogenous CK is able to alleviate SL inhibited bud outgrowth, we used a CK oxidase inhibitor, 1-(2-(2-hydroxyethyl)phenyl)-3-(3-(trifluoromethoxy)phenyl)urea (3TFM-2HE) (Nisler et al., 2021), which reduces degradation of CKs. Like exogenous CK (Dun et al., 2012), 3TFM-2HE treatment promoted additional growth of *rms5* SL-deficient buds (Figure 4D). Similar to other long-term studies with exogenously supplied CKs (Dun et al., 2012), 3TFM-2HE alleviated GR24 inhibited *rms5* bud growth over 7 d (Figure 4D). These results suggest that endogenous CKs act in a similar manner as exogenous CKs and antagonistically with SL to regulate bud outgrowth over long time periods.

### Sucrose and CK can overcome SL inhibited bud outgrowth

Our recent studies have revealed that sucrose can reduce SL response *in vivo* in rice and *in vitro* in pea and rose (Bertheloot et al., 2020; Patil et al., 2021). To test this hypothesis *in vivo* in pea, we examined whether simultaneous treatment of sucrose and SL to intact plants could overcome SL inhibition of bud release. To readily observe SL inhibition, we used SL-deficient plants and supplied sucrose via a syringe to the stem and the synthetic SL, GR24, directly to the measured bud. We found that while GR24 inhibited bud growth of the SL biosynthesis deficient mutant *rms1* (Figure 4E), application of GR24 with sucrose was significantly less inhibitory over the first 2 days (Figure 4E). After 2 days, GR24 was effective at reducing bud growth as described previously in long-term experiments (Dun et al., 2013).

BA completely prevented SL inhibition of bud outgrowth in WT plants over the first 24 h (Figure 4F) (Dun et al., 2012). This lack of SL antagonism of CK response during bud release (shortly after inductive treatments) contrasts with the many findings regarding the antagonism of SL and CK during bud outgrowth which is thought to occur through regulation of *BRC1* (Braun et al., 2012; Dun et al., 2012; Kerr et al., 2020). Hence, we confirmed that this antagonism did indeed occur in the longer term under these experimental conditions (Figure 4F, inset). To test whether a reduced photoassimilate supply may affect the SL/CK antagonism of *BRC1* during bud release, we repeated the experiment under reduced light intensity. When WT plants were grown under lower light conditions, a small but significant antagonistic effect of GR24 and BA was observed in the first 24 h and an antagonistic effect was observed on *BRC1* expression (Supplemental Figure S5). This effect of different light intensities on bud release and *BRC1* expression is consistent with reduced SL signalling under high sucrose conditions (Patil et al., 2021).

Due to the rapid rise in both sucrose and CK content following decapitation (Figure 1) (Mason et al., 2014; Fichtner et al., 2017), the reduced response to SL observed under high sucrose or CK (Figure 4E and 4F) predicts that soon after decapitation, bud growth may be less responsive to SL despite reported effects over the longer term. Indeed, treating axillary buds of WT plants with GR24 prior to decapitation failed to inhibit bud growth within the first 24 h after decapitation (Figure 4G). A significant suppression of bud growth by GR24 in these decapitated plants occurred at 3 days (Figure 3G, inset), which is consistent with the timing used in previous reports of SL-inhibition of bud growth after decapitation in pea (Dun et al., 2013).

### Auxin and GA in buds enhance their sustained outgrowth

The interactions between SL, CK and sucrose have been emphasised above for the early stage of bud growth (bud release; Figure 8). However, to form a branch, the bud must transition to sustained bud growth whereby axillary shoot growth becomes largely independent of the dominance of the main shoot. Many previous studies have explored a role of auxin transport in bud outgrowth and yet in pea, there is little evidence for a role of auxin transport during bud release (Brewer et al., 2015; Chabikwa et al., 2019). As well-established for stem elongation of the main shoot (Yang et al., 1996; O’Neill and Ross, 2002), we also expect an important role of auxin and GA in regulating sustained growth of axillary shoots. To determine if and how GA regulates bud outgrowth in pea, we examined the responses of WT non-growing buds (dormant buds of intact plants) and released buds (activated by CK treatment or decapitation) to GA treatment (Figure 5A and B). Exogenous GA treatments alone did not trigger bud release at any time point in the 3 days following GA treatment (Figure 5A and B). Consistent with a role of GA in sustained bud growth, GA promoted growth of axillary buds released by decapitation or CK at 3 and 5 days after treatment, respectively (Figure 5A and B). The effect of GA on sustained bud growth was further tested by measuring the response to decapitation in a GA biosynthesis deficient mutant of pea (*le*). No significant difference was observed in bud growth between the dwarf *le* and WT plants until day 3 after decapitation when the buds of *le* plants grew significantly less than WT (Figure 5C). Interestingly, GR24 was able to reduce GA-promoted sustained bud growth (Figure 5B).

**Figure 5.**
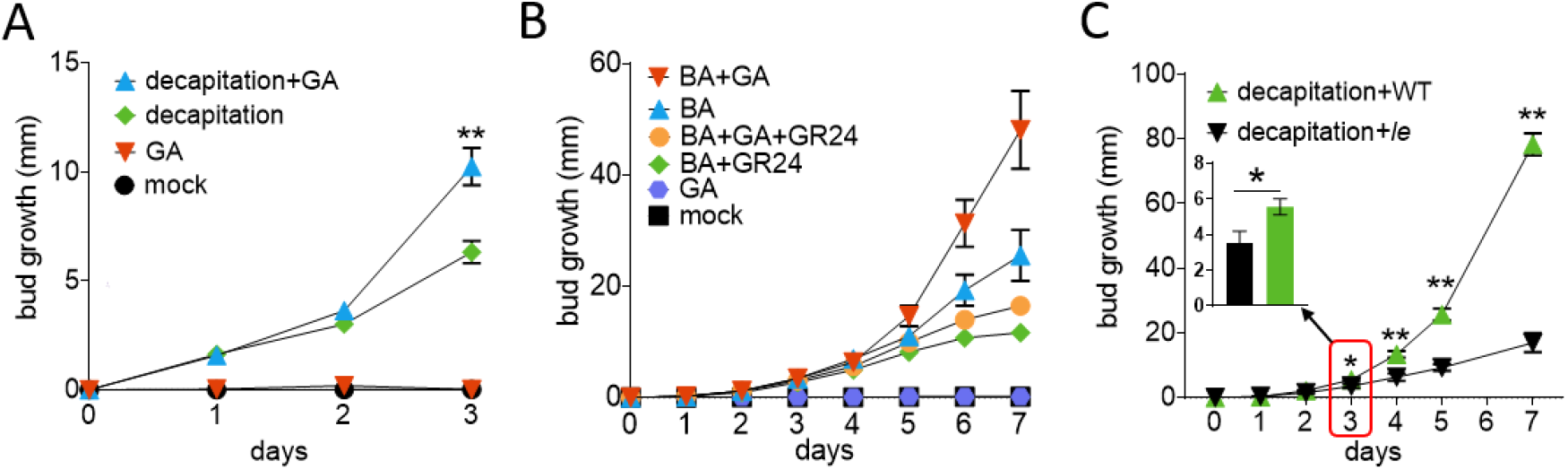
GA does not promote bud release, but rather enhances sustained bud growth. (A) Growth of wild-type node 2 buds after decapitation and/or treatment with 100 μM GA_3_. *n* = 12. ** indicates significant difference between decapitation+GA and decapitation treatment groups. (B) Growth of node 2 wild-type (WT) buds treated with solution containing 0 (mock) or 1g/L GA_3_, and/or 50 μM BA (synthetic CK), and/or 2 μM GR24 (synthetic SL). *n* = 16. (C) Growth of node 4 buds of WT or GA deficient mutant (*le*) plants after decapitation. *n* = 6. All values are mean ± SE. * P <0.05; ** P < 0.01, two-tailed Student’s *t* test.

**Figure 6.**
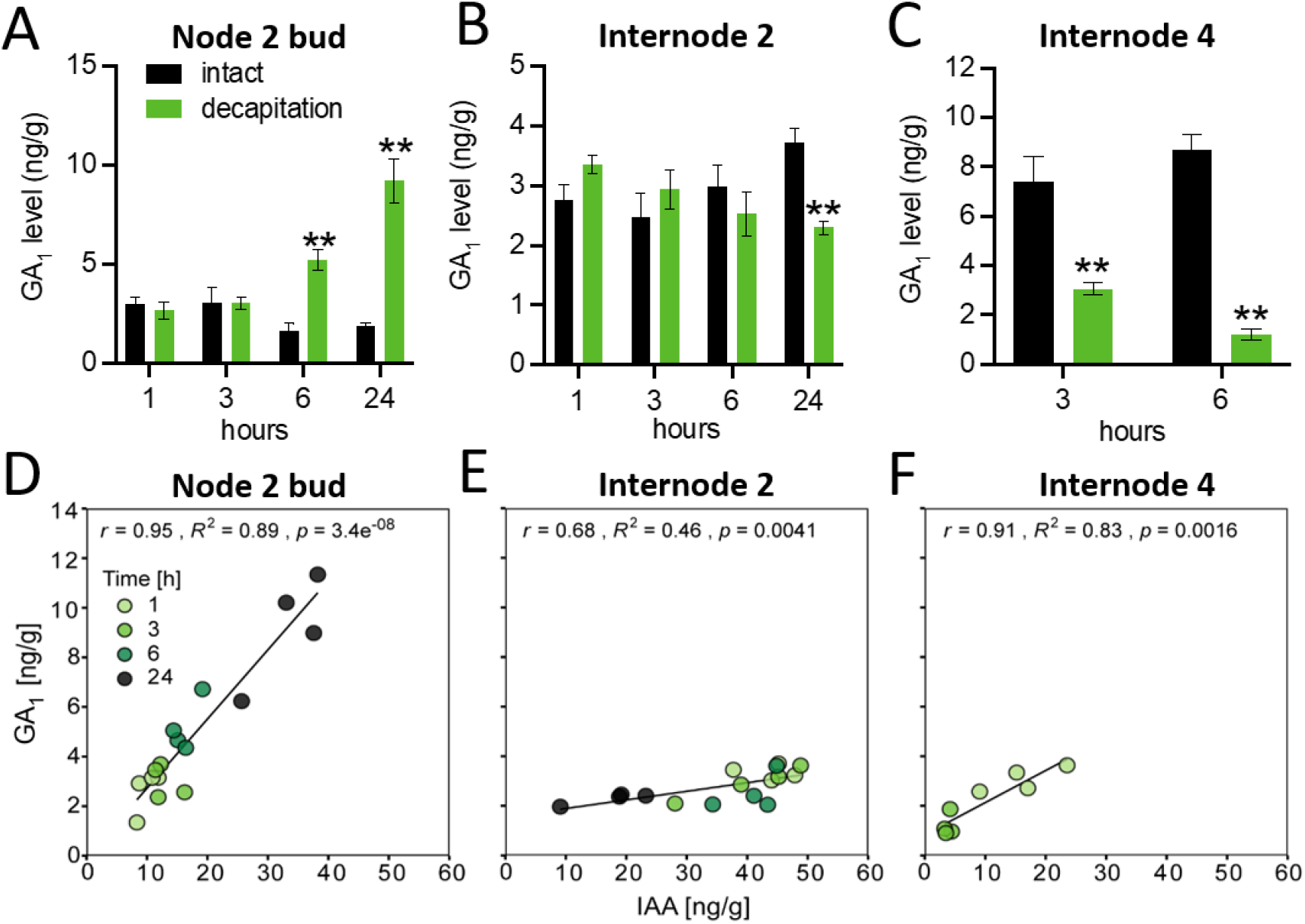
GA level is correlated with auxin level in axillary buds after decapitation. (A, B and C) Endogenous level change of GA_1_ in node 2 bud (A), internode 2 stem (B) and internode 4 stem (C) after decapitation. *n* = 4. Each replicate contains 20 individual buds. Values are mean ± SE, * *P* <0.05; ** *P* < 0.01; Student’s *t* test. (D, E and F) The correlations between GA_1_ and IAA level changes in node 2 buds (D), internode 2 (E) and internode 4 (F). The Pearson correlation coefficient (r), coefficient of determination (R^2^) and probability (*p*) values for each relationship are indicated. These results are from the same plants as in Figure 1.

To establish whether endogenous GA levels may be modulated to affect bud outgrowth, we compared the timing of changes in endogenous GA levels with bud growth in response to decapitation as described in Figure 1. In node 2 buds, levels of GA_1_, the bioactive form of GA in pea, GA_20_ (the precursor to GA_1_) and GA_29_ (a metabolite of GA_1_) significantly increased at 6 h post decapitation (Figure 6A and Supplemental Figure S6A) which is after initial bud growth (2 h; Figure 1). Unlike CK or SL (Braun et al., 2012; Wang et al., 2019), GA treatment had no significant effect on expression of the bud dormancy marker genes, *DRM1*, or *BRC1*, at 6 h after treatment (Supplemental Figure S6B). Combined with the phenotypic responses to GA (Figure 5A-C), these results indicate that GA increases sustained bud growth but has little or no effect on promoting bud release.

Given the known regulation of GA levels by auxin in decapitated plants (Ross et al., 2000; Wolbang and Ross, 2001; Ross et al., 2003), GA_1_ levels in the stem initially decreased within the zone of auxin depletion, but not at the stem below this zone. GA_1_ levels decreased near the decapitation site at 3 h, but only decreased after 24 h in the stem just above node 2, which was positively correlated with auxin level changes (Figure 6B, 6C, 6E and 6F). Consequently, the increase in GA level in node 2 buds at 6 h after decapitation was not correlated with a change in GA or IAA level in the adjacent stem (Figure 6A and 6B, 1C). However, GA and IAA levels in the buds were indeed correlated (Figure 6A and D). Interestingly, this correlation of IAA level and GA level in node 2 buds coupled with the observed effect of GA on sustained growth after bud release, indicates that auxin may act to regulate GA level in growing buds and that GA may act downstream of auxin in promoting sustained bud growth.

Inhibition of auxin signalling, biosynthesis, or efflux out of buds does not affect bud release in pea (Brewer et al., 2009; Brewer et al., 2015; Chabikwa et al., 2019). Here we used decapitated plants with a combined treatment to buds of the auxin perception inhibitor (*p*-chlorophenoxyisobutyric acid; PCIB) and auxin biosynthesis inhibitor (L-Kynurenine; Kyn) and again observed no inhibitory effect on bud release but did observe a significant inhibitory effect on subsequent growth from day 3 (Brewer et al., 2009; Chabikwa et al., 2019) (Figure 7). Consistent with GA action downstream of IAA during this sustained bud growth period, exogenous GA could restore growth to decapitated controls when supplied together with auxin inhibitors (Figure 7).

**Figure 7.**
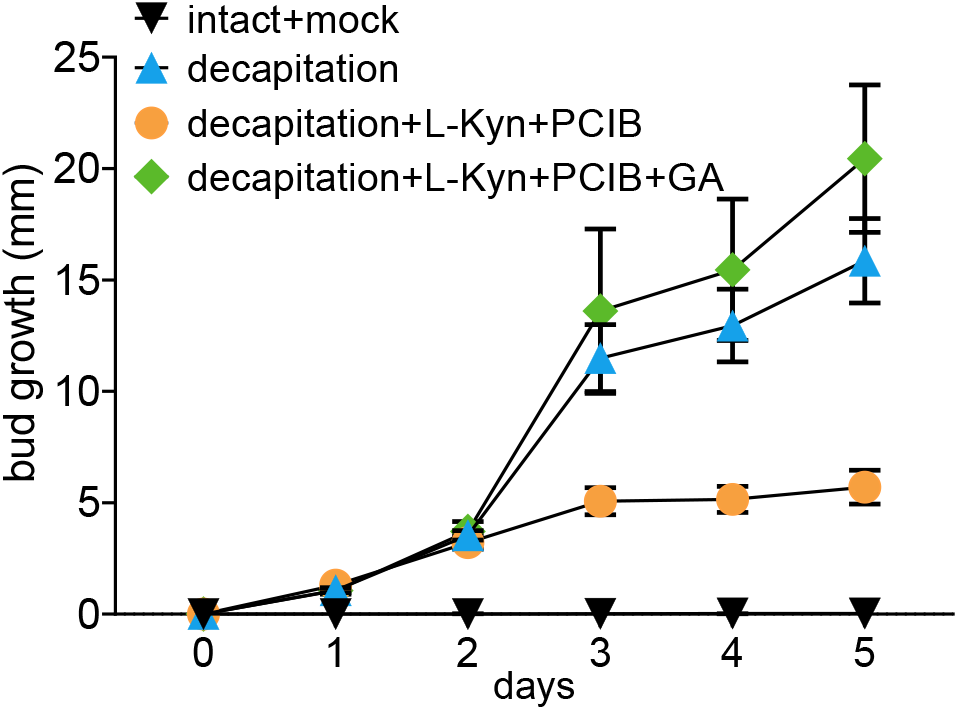
GA can restore decapitation-induced bud growth in absence of auxin. Growth at node 2 after wild-type plants were left intact or decapitated and the buds at node 2 were treated with 10 μl solution containing 0 or 2.5 mM L-Kyn (auxin biosynthesis inhibitor) and 2.5 mM PCIB (auxin perception inhibitor) and/or 500 μM GA_3_. *n* = 6.

## Discussion

### CK and sugars initiate bud release, without stem auxin depletion

By investigating bud outgrowth that occurs outside the zone of auxin depletion after decapitation, we have revealed shortcomings of the classical auxin-centric apical dominance model where auxin depletion after decapitation promotes branching through enhancing CK levels (Sachs and Thimann, 1964; Turnbull et al., 1997; Tanaka et al., 2006). Here we show that changes in stem and bud CK levels following decapitation are not likely due to changes in auxin levels, at least not initially. Auxin depletion occurs too slow to account for the rapid increases observed in CK levels in the stem and bud (Figure 1). CK levels in node 2 buds increased significantly within 1 h of decapitation and before measurable outgrowth or changes in *BRC1* gene expression (Figure 1). These findings demonstrate that decapitation-induced auxin depletion is not the initial signal that triggers CK accumulation in the pea stem and bud distal to the decapitation site (Figure 2). Instead, as suggested previously for stimulating bud outgrowth (Mason et al., 2014), sugars are a strong candidate for this enhancement in CK that occurs outside the zone of auxin depletion (Figure 3).

Sucrose and the sugar signalling metabolite Tre6P accumulate rapidly after decapitation in pea (Mason et al., 2014; Barbier et al., 2015; Fichtner et al., 2017; Fichtner et al., 2021). In rose *in vitro* and dark grown potato, sucrose has been suggested to promote bud outgrowth through enhancement of CK levels (Barbier et al., 2015; Roman et al., 2016; Salam et al., 2021). We used an *in vitro* system to test whether sucrose may enhance CK levels in pea buds. Exogenous sucrose supplied *in vitro* led to somewhat similar changes in CK types to those observed in decapitated plants (Figure 1D and 3B). In buds of sucrose-treated isolated segments and decapitated plants, the levels of *t*Z- and *t*ZR-type CKs consistently increased over time while the accumulation of iP-type CKs in buds stopped at 3-6 h and decreased afterwards (Figure 3B and Supplemental Figure S1C). This supports the premise that rapid enhancement of sucrose levels after decapitation is at least partly responsible for the elevated CK levels (Mason et al., 2014; Fichtner et al., 2017) (Figure 8).

**Figure 8.**
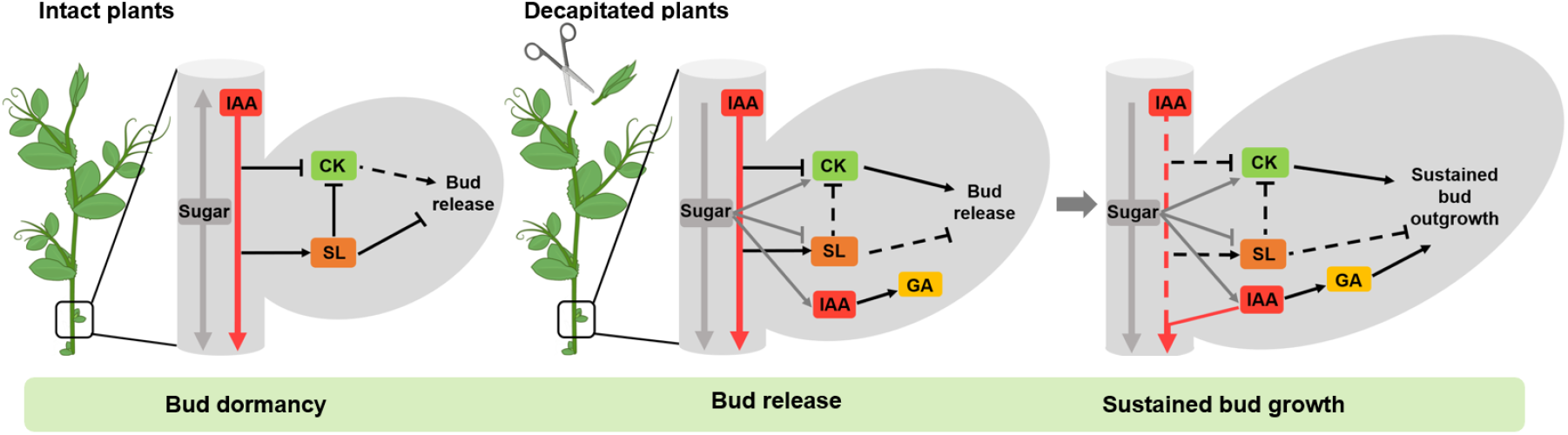
Hypothesis of the network of phytohormone and sugar regulation in apical dominance highlighting different stages including bud dormancy, bud release and sustained bud growth. Dormant buds have very suppressed growth due to the main shoot tip producing auxin and attracting sucrose through its sink strength. This causes comparatively low CK and high SL levels in the stem and buds. After shoot tip removal, rapid accumulation of sugars and CK and reduced SL response trigger bud release. IAA levels in buds also increase at this time consistent with enhanced sugar signalling (Barbier et al., 2015; Ljung et al., 2015). Sustained growth is promoted by continued sucrose supply, together with auxin depletion in the adjacent stem which also enhances CK levels and auxin flow out of buds and reduces SL levels. Enhanced auxin levels in buds promotes GA leading to enhanced bud sink strength and sustained bud growth. The dashed lines indicates a diminished role or effect; flat line inhibition; arrow promotion.

The hypothesis that sucrose may at least in part induce branching through CKs is further supported by the inhibition of sucrose-induced bud growth by inhibitors of CK synthesis or CK perception in potato (Salam et al., 2021). It is also likely that CK increases sugar availability in buds (Ljung et al., 2015; Salam et al., 2021). The recent study in potato suggests a positive feed-forward model whereby sucrose supply to buds enhances CK levels which promotes bud invertase activity, causing a higher bud sink strength which attracts even more sucrose (Salam et al., 2021). This is consistent with the observation that the combined supply of sucrose (up to 50 mM) and exogenous CK (BA) *in vitro* further enhanced the promotion of bud growth in pea (Figure 3C). Moreover, endogenous CK accumulation is likely to have an important effect in pea as chemically reducing endogenous CK degradation, at least in SL deficient buds, greatly enhanced bud growth (Figure 4D).

In decapitated plants, we therefore propose that before stem auxin depletion, rapidly accumulated sucrose and CK act in a module to promote rapid bud release (Figure 8). We propose that, after decapitation, rapid enhancement of sucrose levels in buds followed by the slower depletion of auxin levels in stems promote CK levels over the short and longer term (Figure 2A and 3A) (Schaller et al., 2015). Apical dominance has long been a cornerstone example of the antagonistic relationship between auxin and CK. This study questions the extent to which shoot CK levels are controlled by auxin relative to sugars, and potentially other nutrients (Yoneyama et al., 2020) and reveals a need for future studies on the regulation of CK homeostasis.

### Sugar and CK can over-ride SL signalling during bud release

We used the physiological contexts of decapitation and light quantity to investigate sugar, CK and SL interactions. Decapitation rapidly induces sugar and CK accumulation in buds (Figure 1) (Mason et al., 2014; Fichtner et al., 2017). Using the same GR24 treatment that inhibits bud growth in SL biosynthesis deficient mutants (Figure 4A), there was no significant effect of GR24 on initial decapitation or CK-induced bud growth (Figure 4F and G) despite GR24 being inhibitory after a few days. Similarly, sucrose treatment in intact SL-deficient branching mutants diminished bud inhibition by GR24 within the first two days after treatment (Figure 4E). GR24 treatment under reduced light, and therefore reduced sugar availability, enhanced *BRC1* expression and inhibition of bud release compared with control light conditions (Supplemental Figure S5). These results suggest that the rapid increase in sugar availability (Mason et al., 2014) and CK levels after decapitation (Figure 1) can substantially antagonise the inhibitory effect of SL. This is consistent with the recent findings that sucrose and CK regulate the SL response and/or components of the SL signalling pathway in diverse species including pea, rice and rose (Barbier et al., 2015; Bertheloot et al., 2020; Kerr et al., 2021; Patil et al., 2021). The small size of dormant axillary buds greatly limits their sink strength and ability to attract and utilise sugars. Sugar signalling independently or via CK during bud release (Figure 8) (Barbier et al., 2021) induces buds to grow. This promotion of very small buds may have selective advantage through enabling growth under favourable conditions whilst enabling subsequent inhibition including via competition among growing shoots (Stafstrom, 1995; Barbier et al., 2019; Luo et al., 2021). Future studies should explore sugar fluxes and allocation (Fichtner et al., 2021; Fichtner and Lunn, 2021) that occur during the transition of a bud with high demand for assimilates to a branch comprised of source leaves and an actively growing apical sink. This will provide an excellent context upon which to evaluate the relative contributions of sugar and hormone signalling.

In addition to interactions with sugar pathways, phytohormones also interact with each other to modulate bud release (Wang et al., 2018; Barbier et al., 2019; Luo et al., 2021). Here we demonstrate that exogenous SL treatment in SL-deficient mutants causes a rapid decrease of bud growth and CK levels in axillary buds 6 h after treatment (Figure 4A and B). Consistently, this SL treatment significantly inhibited the expression of CK biosynthesis genes (*IPT1* and *LOG1, 3, 4* and *6*), and promoted the expression of CK catabolism genes (*CKX 3, 5*, and *7*) (Figure 4C). Similar results have been found in peach, where SL treatment on buds decreased decapitation-induced CK accumulation and expression of *IPT* genes (Li et al., 2018). In rice, expression of a CK catabolism gene (*OsCKX9*) was rapidly enhanced by SL treatment and tiller number was enhanced in the corresponding *ckx9* mutant (Duan et al., 2019).

Despite the effect of GR24 on endogenous CK levels, enhanced CK levels are not observed in various shoot tissues of SL-deficient mutants in pea or arabidopsis perhaps due to homeostatic regulation of CK levels over the long-term (Beveridge et al., 1997; Foo et al., 2007; Kiba et al., 2013; Young et al., 2014; Nguyen et al., 2021). In contrast, SL mutants in pea and arabidopsis have greatly suppressed levels of CKs in the xylem sap due to an unidentified systemic shoot-derived feedback signal (Beveridge et al., 1997; Foo et al., 2007). Future studies could assess to what extent the reduction in bud growth by SL is due to independent SL and CK regulation of *SMXL7* and *BRC1* (Dun et al., 2012; Kerr et al., 2021; Patil et al., 2021) versus SL regulation of CK metabolism (Figure 4) (Li et al., 2018; Duan et al., 2019) and whether the systemic shoot-derived feedback signal is related to sugars.

### GA promotes sustained bud growth in pea

After release, buds need to undergo sustained growth to complete their development into branches (Figure 8). By treating released axillary buds with GA (Figure 5), we found that GA promoted sustained bud growth, even though it had no effect on bud release when treated independently on intact plants. A role of GA specific to enhancement of sustained bud growth was supported by the observation that the bud growth difference between GA deficient mutants compared with WT did not occur over the first few days after decapitation (Figure 5C). Moreover, increased endogenous GA levels in buds of decapitated plants did not precede measurable bud release and GA application did not regulate expression of *BRC1* or *DRM1* (Figure 6A, Supplemental Figure S6B). All these results indicate that GA can promote sustained bud growth in pea once buds are released but is itself unable to activate bud release and associated changes in *BRC1* gene expression. It is tempting to speculate that the different effects of GA treatment on buds in different species may relate to whether buds have already entered an initial bud release phase (O’Neill et al., 2019).

In pea shoots, endogenous active GA and auxin levels are well correlated (O’Neill and Ross, 2002; Hedden and Thomas, 2012). As reported previously (Balla et al., 2002), endogenous auxin levels increased in released buds over time (Figure 1C) and is consistent with previous reports that enhanced sugars promote auxin biosynthesis and export from buds during bud release (Sairanen et al., 2012; Barbier et al., 2015). The regulation of sustained growth by GA appears to be tied to the previous established model of auxin- and GA-dependent growth of the main shoot (O’Neill and Ross, 2002; Hedden and Thomas, 2012). Our analysis also revealed a strong correlation between endogenous IAA and GA_1_ levels in internode stems and axillary buds (Figure 6D-F). These results indicate that auxin in buds may induce GA biosynthesis thus prompting sustained bud growth. Auxin efflux from axillary buds can inhibit sustained bud growth but not bud release indicating a role of auxin in buds at an advanced rather than early stage (Brewer et al., 2009; Chabikwa et al., 2019). Moreover, GA can rescue inhibition of sustained bud growth by auxin inhibitors (Figure 7). This also indicates that GA acts downstream of auxin to promote sustained bud growth in pea and provides an alternative suggestion in pea for the auxin transport theory of shoot branching.

### Conclusion and revised model of apical dominance

In the revised apical dominance model (Figure 8), sugars and CK act in a module driving bud release and which suppress inhibition by SL (Figure 1, 3 and 4) (Patil et al., 2021). After an initial bud release stage of growth, SL acts antagonistically against sucrose and CK to suppress subsequent growth (Figure 4) (Dun et al., 2013). Auxin accumulation in released buds promotes sustained growth into branches at least partly through stimulation of GA (Figures 7–8).

## Materials and Methods

### Plant Material, Growth Conditions, Treatments

The lines of garden pea (*Pisum sativum*) used in this study were Torsdag (L107, WT), the GA deficient mutant *le* (NGB5839) and the SL deficient mutant *rms5-3* and *rms1-2T* derived from Torsdag. Plants were grown in 68 mm square pots using the potting mix as previously described (Cao et al., 2020), in a temperature-controlled room (23°C day/18°C night) with an 18-h photoperiod (16 h LED light with 2 h ceiling light extension). Plants with five fully expanded leaves were used unless specified otherwise. Nodes were numbered acropetally from the first scale leaf as node 1 and the distance from node 2 to node 5 was approximately 11 cm. The axillary bud outgrowth at node 2 was monitored following treatments with various combinations of gibberellic acid A3 (GA3), synthetic strigolactone (*rac*-GR24), 6-benzylaminopurine (BA), 3TFM-2HE, L-Kyn and PCIB in 10 μL except for the timelapse experiment with 5 μL. All final solutions contained 1% PEG-1450 and 0.01% Tween-20. The same amount of solvent was added to the control solutions (acetone for GR24, DMSO for 3TFM-2HE, L-Kyn and PCIB and ethanol for BA and GA_3_). For decapitation treatments, the internodes of WT plants were cut 1 cm above node 5. For NAA treatments, 3 g L^-1^ NAA in lanolin was treated at the cut stump immediately after decapitation as described previously (Foo et al., 2005). The sugar treatment *in vivo* was followed as described previously (Mason et al., 2014).

### Phenotypic analysis of bud outgrowth

The measurement of bud length was performed using time-lapse photography at 1 h intervals, as described previously (Mason et al., 2014). The daily measurement of bud length was performed using digital callipers (resolution: 0.01 mm). Bud growth was calculated as the difference between the initial and current bud size.

### *In vitro* cultivation of pea axillary buds

Previous established method was used (Barbier et al., 2015). Briefly, node 2 pea stem segments (1.5 cm) were excised from intact plants with five expanded leaves. Stipules and leaves were removed before the stem segments were transferred onto half strength Murashige and Skoog growth medium, supplemented with 50 mM of sucrose or mannitol. The plate containing stem segments were cultured in the growth room as described above.

### Gene expression and phytohormone profiling

Total RNA and phytohormones were extracted and processed as described previously from the same plant materials and using internal standards for the phytohormones (Barbier et al., 2019; Cao et al., 2020). Three to four replicates were used, each containing 10-20 buds. *PsEF1α, PsGADPH* and *PsTUB2* were used as RT-PCR reference genes for normalization. Primer sequences are listed in Supplemental Table S1.

### Data processing and statistical analysis

Data analysis for gene expression and phytohormone profiling was performed as described previously (Cao et al., 2020). Statistical analyses were performed using Graphpad prism 9.0 (Graphpad Software, USA) and correlation analysis was performed using R with Pearson’s correlation. Two tailed Student’s *t*-test and one-way ANOVA (Fisher’s LSD test) were used unless otherwise stated.

## Acknowledgments and Funding

The authors acknowledge the scientific and technical assistance from Dr. Amanda Nouwens and Mr. Peter Josh for mass spectrometry and Dr Jaroslav Nisler for providing the CKX inhibitor. This research was funded by the Chinese Scholarship Council, The University of Queensland, the Australian Research Council (Georgina Sweet Laureate Fellowship, FL180100139; Centre of Excellence for Plant Success in Nature and Agriculture, CE200100015). This work is dedicated to the memory of our good friend and colleague, Tinashe Chabikwa, who died in March of 2022, during the preparation of this paper.

## Author Contributions

All authors contributed to experimental design and critically reviewed the manuscript. D.C., L.D., and S.K. performed experiments and analyzed data. D.C., C.B., F.B., E.D. and F.F. interpreted data. D.C. and C.B. wrote the manuscript with input from the other authors, and all authors edited the manuscript.

## Supplemental Material

**Supplemental Figure S1.** Endogenous CK levels in internode 4 (A), internode 2 (B), node 2 bud (C) at 1, 3, 6, and 24 h after decapitation. Node 2 was about 12 cm from the decapitation site. * P<0.05, **P < 0.01, Student’s t test, n = 4. Each replicate contains 20 individual buds. Values are mean ± SE. Abbreviations: iPRMP, isopentenyladenosine-5’-monophosphate; tZRMP, transzeatin riboside-5’-monophosphate; DZRMP, dihydrozeatin riboside-5’-monophosphate; iPR, isopentenyladenosine; tZR, trans-zeatin riboside; DZR, dihydrozeatin ribodide; iP, isopentenyladenine; tZ, trans-zeatin; DZ, dihydrozeatin; IPT, adenosine phosphate-isopentenyltransferase; LOG, cytokinin phosphoribohydrolase ‘Lonely guy’; CYP735A, cytochrome P450 mono-oxygenases. ND, not detected.

**Supplemental Figure S2.** (A, B) Extracted ion chromatograms showing a detectable NAA signal (186.9->141 m/z transition, 9.8 min retention time) in internode 4 (A) and undetectable NAA signal in internode 2 (B) 4 h after decapitation and treatment with 3g/L NAA applied to the decapitated stump above internode 4. (C, D) Endogenous GA_1_ levels in internode 4 (C) and internode 2 (D) 4 h after decapitation of decapitated plants treated either with mock or 3 g/L NAA above internode 4. Values are mean ± SE, *n* = 4. Multiple comparison tests were performed with one-way ANOVA; *n* = 4.

**Supplemental Figure S3.** Buds of nodal stem segments exhibit growth after 24 h treatment with 50 mM sucrose compared to 50 mM mannitol control.

**Supplemental Figure S4**. (A) Endogenous CK levels in *rms5* node 2 buds and internode 2 treated with or without 10 μM of GR24 on node 2 buds for 6 h. n = 3. (B) Expression of bud dormancy genes in *rms5* node 2 buds treated with or without 10 μM of GR24 for 6 h. *n* = 3. Values are mean ± SE. Each replicate contains 20 individual buds. * P<0.05, **P< 0.01, Student’s t test.

**Supplemental Figure S5.** The early response of CK treated buds to SL is reduced under higher light. Effect of GR24 on BA-induced bud outgrowth at 24 h (A) and *BRC1* expression at 6 h (B) after treatment. Treatments were 50 μM BA ± 5 μM GR24; normal light, 150-200 μmol m^-2^s^-1^; low light, 50-75 μmol m^-2^s^-1^. Expression of *BRC1* in the bud at node 2 is represented relative to the high light mock control. Different letters on the top of columns indicate significant difference with one-way ANOVA. Values are mean ± SE; *n* = 6 plants (A) or 6 pools of 6 plants (B).

**Supplemental Figure S6.** (A) Changes in levels of endogenous GAs (GA_1_, GA_20_, GA_29_) in node 2 bud, internode 2 and internode 4 after decapitation. Each replicate contains 20 individual buds; *n* = 4. Values are mean ± SE. * *P* <.05; ** *P* < 0.01, with respect to the directly comparable treatment; Student’s *t* test. NA, not available. (B) Node 2 axillary buds were treated with 2.9 mM GA or 50 μM BA for 6 hours. Values are mean ± SE, *n =* 4. Each replicate contains 20 individual buds. **P < 0.01 compared to mock control, Student’s *t* test.

**Supplemental table S1.** Primers used in the study.

